# NAPRT expression and epigenetic regulation in pediatric rhabdomyosarcoma as a potential biomarker for NAMPT inhibition

**DOI:** 10.1101/2025.07.28.667284

**Authors:** Angela Kim, Prateek Bhardwaj, Collin D. Heer, Sophia J. Zhao, Karlie N. Lucas, Katelyn J. Noronha, Sam Friedman, Raffaella Morotti, Ranjini K. Sundaram, Shahyan Rehman, Deepti Bhatt, Filemon S. Dela Cruz, Tamar Y. Feinberg, Subhash Ramakrishnan, Wei Xue, Donald A. Barkauskas, David Hall, Jack F. Shern, Wenyue Sun, Frederic G. Barr, Jae-Sung Yi, Josh Spurrier, Amy Yu, Charles Brenner, Christine M. Heske, Juan C. Vasquez

## Abstract

**Purpose:** New treatments are needed to improve survival in children with rhabdomyosarcoma (RMS). NAD⁺ biosynthesis, regulated by the enzymes NAPRT and NAMPT, represents a metabolic vulnerability due to high NAD⁺ turnover in cancers. Although NAMPT inhibitors (NAMPTi) show preclinical promise, clinical translation has been limited by toxicity and the lack of predictive biomarkers. Here, we evaluated NAPRT expression in RMS and its potential as an actionable biomarker to guide NAMPTi therapy.

**Experimental Design:** NAPRT promoter methylation, transcript levels, and protein expression were assessed in RMS cells, PDXs, and primary tumors (n=109) from the Children’s Oncology Group. *In vitro* sensitivity to NAMPTi was tested in molecularly diverse and isogenic RMS cell lines, examining the role of NAPRT expression in mediating cytotoxicity and the ability of nicotinic acid (NA) to rescue viability. *In vivo* efficacy was assessed using NAPRT-isogenic orthotopic xenograft models.

**Results:** NAPRT promoter hypermethylation was found in a subset of RMS models and patient samples. Immunohistochemistry showed loss of NAPRT protein in 30–40% of tumors, defined as <1% tumor cell staining. Methylation modestly correlated with protein expression. NAPRT-silenced cells were highly sensitive to NAMPTi, driven by NAD⁺ depletion and not reversible with NA. *In vivo,* NAMPTi induced significant tumor regression, which was not abrogated with NA administration in NAPRT-silenced models.

**Conclusions:** NAPRT loss occurs in a subset of RMS, offering a potential strategy to expand the therapeutic window of NAMPTi. Further research is needed to understand NAPRT regulation and optimize biomarker assay strategies for use in future clinical trials.

## Introduction

Rhabdomyosarcoma (RMS) is the most common soft tissue sarcoma among children, accounting for 5 percent of all pediatric cancers (1). The current treatment strategy for RMS consists of chemotherapy, surgical resection, and/or radiation therapy. While outcomes for patients with low-risk RMS are excellent, the long-term survival rates for patients with metastatic or high-risk disease is less than 30 percent (2). As there have been no significant improvements in the overall survival for pediatric patients with high-risk RMS over the last several decades, there is a dire need to develop new treatments (3,4).

Nicotinamide adenine dinucleotide (NAD^+^) plays a vital role in cellular energy metabolism as an essential coenzyme for glycolysis, the tricarboxylic acid (TCA) cycle, oxidative phosphorylation, fatty acid metabolism, antioxidant metabolism, and serine biosynthesis, which all contribute to cancer progression (5–7). Furthermore, cancer cells exhibit several characteristic features that drive their reliance on NAD^+^ production, including dysregulated metabolism, rapid proliferation, and continuous NAD^+^ consumption by NAD^+^-degrading enzymes, such as PARPs and sirtuins (8,9).

Given this dependency, numerous studies have suggested that disrupting NAD^+^ biosynthesis could serve as a potential therapeutic strategy (10). Cells synthesize NAD^+^ primarily through the Preiss-Handler and Salvage pathways driven by nicotinic acid phosphoribosyltransferase (NAPRT) and nicotinamide phosphoribosyltransferase (NAMPT), respectively **(Figure 1A)** (10). Preclinical studies have demonstrated that pharmacological NAMPT inhibition effectively depletes NAD^+^, leading to cytotoxicity in cancer cells due to their increased metabolic demand (11–14). However, early-phase clinical trials of NAMPT inhibitors (NAMPTi) in unselected adult patients with advanced cancers showed few objective responses and dose-limiting toxicities including bone marrow suppression, particularly thrombocytopenia (15–17). While retinal and cardiac toxicities were noted in pre-clinical animal studies, this was not reported in humans (18–20).

**Figure 1.**
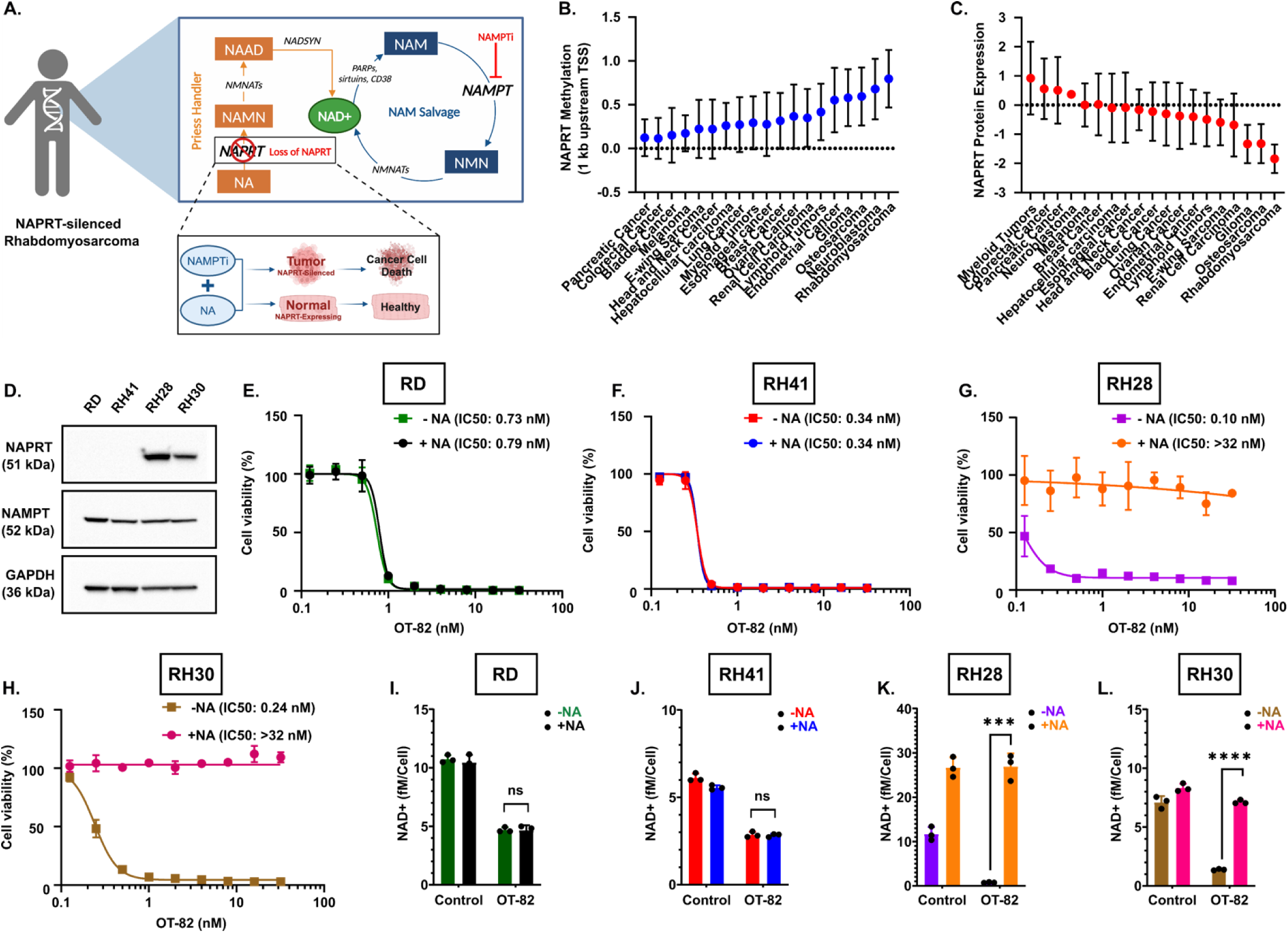
RMS models display loss of NAPRT and sensitivity to OT-82 that is not rescued through the Preiss-Handler pathway. **A.** Schematic of NAD^+^ biosynthesis pathways and “protective rescue” paradigm in tumors with loss of NAPRT. **B.** Plot illustrating the methylation of NAPRT across various cancer types. NAPRT is hypermethylated in RMS. **C.** Plot illustrating NAPRT protein expression across various cancer types. Loss of NAPRT protein expression is observed in RMS. **D.** Protein expression of NAPRT in four RMS models by western blot. **E-G.** Cell viability in NAPRT-silenced RD (**E)** and RH41 (**F)** cells treated with increasing concentrations of OT-82 for 6 days with or without 10 μM NA. **G-H.** Cell viability in NAPRT-expressing RH28 (**G**) and RH30 (**H**) treated with increasing concentrations of OT-82 for 6 days with or without 10 μM NA. **I-L.** Total NAD^+^ quantification post-24 hours in RD (**I),** RH41 (**J**), RH28 (**K**) and RH30 (**L**) treated with respective IC50 concentrations with or without 10 μM NA. The data is plotted as mean with error bars indicating SEM.*p<0.05; ***p<0.001; ****p<0.0001; n.s., not significant. NAPRT, nicotinate phosphoribosyltransferase; NAMPT, nicotinamide phosphoribosyltransferase; NA, nicotinic acid; GAPDH, Glyceraldyde3phosphate dehydrogenase.

NAMPTi have not been clinically tested in pediatric solid tumors. Notably, there is emerging evidence that pediatric bone and soft-tissue sarcomas show exquisite sensitivity, potentially due to unique metabolic vulnerabilities and intrinsic defects in DNA damage repair that are exacerbated by inhibition of NAD^+^-dependent DNA repair enzymes (9,21–27). However, the lack of a known actionable biomarker still limits clinical translation. Most normal tissues express NAPRT while a significant proportion of tumor tissues do not (13,28). Previous work from our group and others indicates that loss of NAPRT expression by tumor-specific promoter CpG island methylation may provide a novel biomarker for sensitivity to NAMPTis (9,13,26,28–31).

In this study, we found that a subset of tumors across multiple RMS subtypes, including both FOXO1 fusion–positive and –negative, as well as embryonal and alveolar histologies, harbor NAPRT promoter methylation and loss of NAPRT protein expression. Moreover, NAPRT-silenced RMS models were highly sensitive to NAMPTi *in vitro* and cytotoxicity was not rescued by nicotinic acid (NA) supplementation, suggesting that the Preiss-Handler pathway is not functional in these cells (**Figure 1A**). *In vivo,* NAMPTi resulted in significant tumor regression in NAPRT-silenced RMS orthotopic models, which was not reversed by NA administration, whereas NA completely abrogated NAMPTi efficacy in NAPRT-expressing tumors. These observations were emulated by rescue of NAD^+^ pools only in NAPRT-expressing models upon NA supplementation. Overall, these data suggest that NAPRT loss may serve as a therapeutic target in RMS tumors by inducing a synthetic lethal interaction with NAD^+^ depleting agents and providing an opportunity to widen the therapeutic index with NA supplementation.

## Materials and Methods

### Cell Culture

RD cells were cultured in Dulbecco’s Modified Eagle Medium (DMEM) (Gibco). RH28, RH30, and RH41 cells were cultured in Roswell Park Memorial Institute (RPMI) (Gibco). All media formulations contained 10% fetal bovine serum (FBS) (Sigma) and 1% penicillin-streptomycin (Gibco). Mycoplasma testing was performed routinely using the MycoAlert assay kit (Lonza Biosciences, Durham, NC, USA). All cell cultures were maintained and treated in 5% CO2, 20% O2, and humidified in a 37°C incubator unless otherwise noted. Cells were collected for experiments 1 to 8 weeks after thawing. Cell lines were authenticated through the Yale Keck Biotechnology Core with short tandem repeat (STR) genotyping by fragment analysis. STR profiles generated via the Yale Keck Core will be made available upon request.

### Isogenic Model Generation

RH41 cells were transfected with 2 μg of NAPRT cDNA plasmid (Genscript, Cat. # OHu28558D) using Lipofectamine 3000 (ThermoFisher). 48 hours post-transfection, cells were treated with G418 (Geneticin) to select for NAPRT-expressing cells. When cells were ∼80% confluent, a flow-sorted single cell generation was performed in 96-well plates. NAPRT overexpression was confirmed through immunoblot and functionally validated through cell proliferation assay.

RH30 cells were transfected with 37.5 pM of respective sgRNA (CRISPR948669_SGM (gRNA1); CRISPR948672_SGM (gRNA2)) and 6.25 μg of Cas9 enzyme using Lipofectamine CRISPRMAX (ThermoFisher Scientific). When cells were 70∼80% confluent, approximately 48 hours post-transfection, single-cell clones were isolated using a limited dilution cloning in 96-well plates. NAPRT KO was validated by immunoblotting and functional cell-based assay.

### In Vitro Chemical Treatments

FK866 (Selleckchem) and OT-82 (Selleckchem) were dissolved in DMSO and used for treatment as indicated. NA (Sigma) was solubilized in 1M NaOH, then diluted in complete media immediately prior to treatment alone or in combination with FK866 or OT-82, as indicated. NMN (Sigma) was diluted in PBS prior to treatment. NR (SelleckChem) was diluted in DMSO prior to treatment. NA, NMN, and NR were aliquoted and stored at −20°C.

### Cell Viability Assays

Cells were plated at a density of 1,000 to 8,000 cells per well in 96-well plates and treated with NAMPTi 24 hours after plating. For rescue experiments, 10 μM of NA was simultaneously administered with NAMPTi to cells during the treatment phase. Following 4 to 6 days of incubation, cells were fixed with 4% paraformaldehyde and stained with 1µg/ml Hoescht 3342. Cells were imaged with a Cytation3 (BioTek) and counted via CellProfiler (https://cellprofiler.org, RRID:SCR_007358). Each experiment was conducted with at least three technical and biological replicates. Data was normalized to the vehicle-treated control group prior to analysis in GraphPad Prism Software with a non-linear regression analysis “inhibitor vs response – variable slope (four parameters).”

### PDX IncuCyte Analysis

Real-time longitudinal cell proliferation was measured using the IncuCyte SX5 live-cell analysis system (Sartorius) with cell density and morphology imaged every 8 hours. PDX-derived RMS cell line SJRHB013758 (FOXO1 fusion-negative) was plated at a density of 2000 cells/well in 96-well plates. Cell culture conditions for this model have been previously described here (25). Cells were allowed to adhere overnight and treated the following day with doses of OT-82 (OncoTartis), nicotinic acid (Sigma) at 10 μM, or both. Each experiment was conducted 3 times with 12 technical replicates.

### Apoptosis Assays

Cells were plated at a density of 1.5 x 10^5^ cells per well in 6-well plates and treated with DMSO, NA, NAMPTi, or concurrent NA and NAMPTi 24 hours after plating. After 48 hours of treatment, Annexin V/PI apoptosis assay was performed using a Dead Cell Apoptosis Kit (ThermoFisher) as per manufacturer’s specification. Annexin V/PI assay was performed in three technical replicates.

### Cell Cycle Analysis

Cells were plated at a density of 1.5 x 10^5^ cells per well in 6-well plates and treated with DMSO, NA, NAMPTi, or concurrent NA and NAMPTi 24 hours after plating. After 48 hours of treatment, cells were trypsinized and fixed dropwise with 70% ice-cold ethanol. Following, cells were incubated with Propidium Iodide/RNase A stain (BD Biosciences) for 15 minutes at room temperature and analyzed through flow cytometry.

### Western Blotting

Protein (40μg/lane) was separated by 15% SDS-PAGE (Bio-Rad) and transferred onto polyvinylidene difluoride membranes (Immobilon). Membranes were blocked with 5% nonfat skim milk for 1 hour at room temperature then incubated with primary antibodies at 4°C overnight (**Supplemental Table 1**). Membranes were washed in TBST (20 mM Tris pH 7.5, 150 mM NaCl, 0.1% Tween20) and then incubated with secondary antibodies (Proteintech and Cell Signaling Technology) for 1.5 hours at room temperature. Signals were visualized with an ECL Kit (Bio-Rad).

### NAD^+^/NADH Quantification

Cells were plated at 4,000 cells/well in 96-well plates and treated 24 hours later with NAMPTi with and without NA concurrent supplementation for 24 hours. NAD/NADH-Glo Assay (Promega) was performed per the manufacturer’s specification. NAD/NADH-Glo Assay was performed in three biological replicates.

### In Vivo Studies (efficacy and NAD^+^ quantification)

Animal studies were carried out in accordance with Yale University’s Institutional Animal Care and Use Committee guidelines. Four-to six-week-old female Fox Chase SCID Beige mice (CB17.B6-PrkdcscidLyst bg/Cr) from Charles River Laboratories (n = 3-5 mice per treatment group) were used for cell line xenograft experiments. Two million cells were suspended in PBS and matrigel at a final volume of 100 µl and injected into the gastrocnemius muscle. Mice were randomized once the tumor was palpable. Mice were treated with vehicle (30% cyclodextrin), OT-82 (25 mg/kg), or OT-82 in combination with NA (25 mg/kg) by oral gavage daily, following a 3-days on and 4-days off treatment cycle for 5 weeks. Tumors were measured twice/week with calipers and volume was calculated by V = 0.5 x L x W^2^.

For *in vivo* NAD^+^ quantification, tumors and normal muscle were harvested from the mice 2 hours after 1 treatment cycle (i.e., after 3 days ON period of NAMPTi dosing). All tissues were snap-frozen. NAD/NADH-Glo Assay (Promega) was performed per the manufacturer’s specification. NAD/NADH-Glo Assay was performed in three technical replicates with 3-4 biological replicates per treatment group (Vehicle, NAMPTi, NAMPTi+NA). NAD^+^ concentration was normalized to the total mass of tissue (mg).

### PDX tumor tissue microarray

Patient-derived xenograft (PDX) tissue microarrays (TMAs) were constructed using formalin-fixed and paraffin-embedded PDX tissues. Paraffin sections from 29 pediatric RMS PDX models were stained with Hematoxylin and Eosin (H&E) to obtain a template guide slide for each tissue block and to validate sample quality and preservation. PDX samples with greater than 20% necrosis or inadequate tissue preservation were excluded. Slides were annotated to create a template for punching and used as guides for PDX tumor core selection. Each array was designed using 0.6 mm diameter punches, with duplicates for each model. Tumor tissue cores from the 29 RMS PDX models were included in the arrays. Human (tonsil, testis, breast, colon, prostate, lung, kidney, liver) and mouse (spleen, skin, colon, kidney, liver) tissues were also included as controls.

### Rhabdomyosarcoma patient samples

Human tissue procurement and analysis procedures for specimens from patients with soft-tissue sarcoma were approved by the Institutional Review Board (IRB #2000027341) at Yale University. TMA slides containing patient RMS samples were also obtained from the National Cancer Trial Cooperative Network (NCTCN) and the Children’s Oncology Group (COG). Samples included in the TMAs were collected from patients enrolled in COG clinical trials D9602, D9802, D9803, following institutional review board (IRB)-approved protocols. The COG TMAs comprised 278 cores from 130 patients aged 0–18 years of both embryonal (FOXO1 fusion negative), and alveolar (FOXO1 fusion positive) tumors. We excluded patients where only control cores and/or cores with insufficient tissues were presented, resulting in a final analysis of 109 pediatric patients. Archived formalin-fixed paraffin-embedded tissue was used to construct the TMA with cores each measuring 1.0 mm in diameter. Presence of tumor was assessed *via* H&E staining.

### NAPRT immunohistochemistry

The preparation of the TMA tissue for immunohistochemistry (IHC) was performed with an anti-NAPRT mouse monoclonal antibody (4A5D7), developed, and validated by Promab at Alphina therapeutics (9). NAPRT IHC specificity and staining optimization were performed on cell line models and normal human tissue by NeoGenomics Laboratories using a Leica Bond III. Briefly, epitope retrieval was performed for 25 minutes at 100°C, blocked with Leica Protein Block for 30 minutes, and incubated with primary antibody (0.5 µg/mL) for 30 minutes. 3-3-Diaminobenzidine (DAB) was used for colorimetric detection. Samples were counterstained with hematoxylin. NAPRT signal was scored according to the following criteria by a pathologist. The semi-quantitative IHC scoring systems utilized are described as follows. Percentage Positive Score: (Score = % Positive Cells) 0 = <1%; 1+ = 1-25%; 2+ = 25-50%; 3+ = 50-75%; 4+ = 75-100%.

### DNA methylation analysis

Genomic DNA from 29 RMS tumors was analyzed using the Infinium MethylationEPIC BeadChip (Illumina). In addition, DNA methylation data from 70 RMS tumors previously generated on the Infinium HumanMethylation450 BeadChip were included in this analysis (dbGaP: phs001970) (32). Raw IDAT files from both sources were processed and normalized using the swan algorithm in the minfi package (https://bioconductor.org/packages/release/bioc/html/minfi.html) (33,34). The β-value was computed as the measure of methylation, ranging from 0 (completely unmethylated) to 1.0 (completely methylated). The CpG probe annotation corresponding to the NAPRT1 gene was retrieved using the Illumina Human Methylation EPI Canno.ilm10b2.hg19 package (https://bioconductor.org/packages/release/data/annotation/html/IlluminaHumanMethylationEPI Canno.ilm10b2.hg19.html).

### RNA sequencing and expression analysis

RNA sequencing data was available for 31 COG RMS samples as previously described and available in the NCI Oncogenomics expression database (https://omics-oncogenomics.ccr.cancer.gov/cgi-bin/JK) (35,36). Briefly, RNA was extracted from samples using the Qiagen RNeasy Micro Kits according to the manufacturer’s protocol (Qiagen). PolyA selected RNA libraries were prepared for RNA sequencing on Illumina HiSeq2000 using TruSeq v3 chemistry according to the manufacturer’s protocol (Illumina). Hundred bases–long paired-end reads were assessed for quality and reads were mapped using the bcl2fastq tool in CASAVA (Illumina). Fastq files were mapped to GRCH37 reference genome using the STAR alignment algorithm and quantified by the RSEM program based on Ensembl GFCh37.75 gene annotation (37,38). Read counts were normalized using the TMM method in edgeR and transformed to FPKM (39).

### Statistical Analyses

Student two-tailed *t* test was used for comparing two groups. One-way ANOVA with Dunnett’s multiple comparison test was used to evaluate experiments involving one variable across multiple groups. Two-way ANOVA with Tukey’s multiple comparison test was used to evaluate two variables across multiple groups. Log-rank test was performed to compare survival of mice. Statistical significance was defined as p < 0.05. Statistical analyses were carried out using GraphPad Prism 10 software. All experiments were performed in biological triplicates unless noted otherwise.

## Results

### RMS models display NAPRT promoter methylation and low protein expression

We probed data from the Cancer Dependency Map (DepMap) to assess for NAPRT promoter methylation and protein expression across different cancer types (40). High rates of NAPRT promoter methylation accompanied by low levels of NAPRT expression were observed in RMS cell line models **(Figure 1B and C).** We then went on to probe for NAPRT protein expression by immunoblot in a panel of RMS cell lines representing both FOXO1 fusion negative (RD) and FOXO1 fusion positive (RH28, RH30, RH41) disease (41). We confirmed that a subset of models (RD and RH41) do not express NAPRT at the protein level, while NAMPT was detected in all cell lines (**Figure 1D**).

### NAPRT-silenced RMS cells show sensitivity to NAMPTi that is not reversed with nicotinic acid (NA) supplementation

To determine whether NAPRT expression affects sensitivity to NAMPT inhibitors (NAMPTi), RMS cells were treated for 96 hours with NAMPTi, with or without concurrent NA supplementation. RMS cells demonstrated marked sensitivity to FK-866, a first-generation small-molecule NAMPTi, as well as to OT-82, a newer NAMPTi that has shown reduced cardiac, neurological, and retinal toxicities in preclinical studies compared to earlier NAMPT inhibitors tested in clinical trials (15). As expected, cells with a functional Preiss–Handler pathway driven by NAPRT expression (RH28 and RH30) were rescued by NA supplementation, whereas NAPRT-silenced cells (RD and RH41) exhibited no change in viability **(Figure 1E-H, Supplemental Figure S1A-D).**

Next, NAD^+^ levels within RMS cell lines were quantified to better understand the mechanism underlying their sensitivity to NAMPTi. NAD^+^ levels were assessed 24 hours post-treatment with FK-866 or OT-82. Pools of NAD^+^ dropped markedly in NAMPTi-treated cells that did not express NAPRT (RD and RH41) both in the absence and presence of NA, confirming their enhanced dependence on NAMPT activity for survival **(Figure 1I and J, Supplemental Figure S1E** and **F).** In contrast, NA supplementation fully restored NAD^+^ levels after NAMPTi treatment in cells that expressed NAPRT (RH28 and RH30) (**Figure 1K** and **L, Supplemental Figure S1G** and **H)**. A PDX model (SJRHB13758), characterized as NAPRT-expressing via immunoblot, demonstrated *ex vivo* sensitivity to OT-82, with rescue through a functional Preiss-Handler pathway upon co-administration of NA, at concentrations similar to those used in cell line models (**Supplemental Figure S1I** and **J).**

To verify the on-target mechanism of NAMPTi and NAPRT-dependent rescue, we performed *in vitro* cell viability assays with concurrent supplementation of nicotinamide mononucleotide (NMN), nicotinamide riboside (NR), or NA. Nicotinamide is converted to NMN by NAMPT in the classical salvage pathway; therefore, regardless of NAPRT status, we would expect to observe rescue in cell viability (10). Similarly, NR is phosphorylated by nicotinamide riboside kinase (NRK1/2) to form NMN upstream of NAMPT (10). As hypothesized, RH41 (NAPRT-silenced) and RH30 (NAPRT-expressing), were both rescued with co-administration with NMN and NR (**Supplemental Figure S2A,C,D,F)**. However, when concurrently supplementing with NA, only RH30 was rescued following NAMPTi treatment (**Supplemental Figure S2B,E**).

To confirm the dependence on NAPRT expression in mediating NAMPTi response and rescue with NA, NAPRT was exogenously introduced into RH41 cells **(Figure 2A, Supplemental Figure 3A)** and RD cells **(Supplemental Figure 3D).** Ectopic expression of NAPRT in RH41 (RH41^NAPRT+^) and RD cells (RD^NAPRT+^) rescued cell death with co-administration of NA, which emulated the effect of NAMPTi with NA supplementation in cells with endogenous NAPRT expression (RH28 and RH30) **(Figure 2B and C, Supplemental Figure S3B,C,E).** To further corroborate this observation, we performed CRISPR/Cas9-mediated knockout (KO) of NAPRT in naturally expressing RH30 cells (RH30^NAPRT-^). Knockout of NAPRT was functionally validated via immunoblot, and cell proliferation assays demonstrated that RH30^NAPRT-^ mimicked the effect of NAMPTi without NA rescue, as is seen in endogenously NAPRT-silenced cells (**Figure 2E-G, Supplemental Figure S4A-C**). Furthermore, NAD^+^ levels were assessed in these isogenic models following a 24-hour NAMPTi treatment. Both FK866 and OT-82 significantly diminished the NAD^+^ pool in both RH41 wild-type and RH30^NAPRT-^, even with co-administration of NA, while RH41^NAPRT+^ and RH30 control were able to rescue their NAD^+^ levels (**Figure 2D** and **H**). Collectively, these data demonstrate that RMS cell lines are highly sensitive to NAMPT inhibition through NAD^+^ depletion, with NA failing to rescue NAPRT-silenced cells, while NAPRT-expressing cells are protected by NA co-treatment.

**Figure 2.**
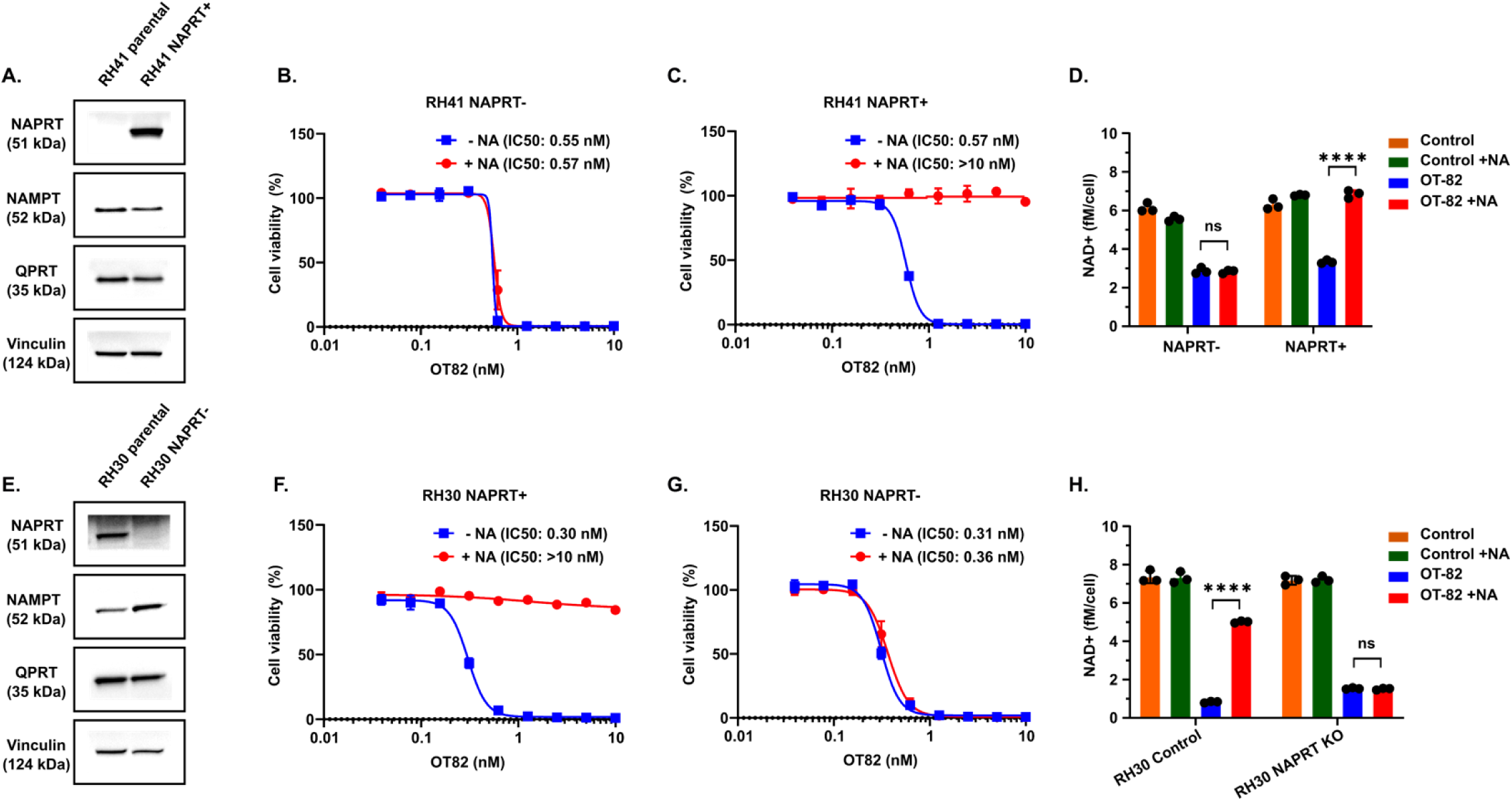
Generation and functional validation of NAPRT isogenic models. A-D. Endogenously NAPRT-silenced RH41 cells transfected to ectopically express NAPRT (RH41^NAPRT+^). **A**. Protein expression of NAPRT, NAMPT, QPRT, and Vinculin in RH41 parental and RH41^NAPRT+^. **B-C.** Cell viability in RH41 (NAPRT-silenced) (**B**) and RH41^NAPRT+^ (**C)** treated with increasing concentrations of OT-82 for 4 days with or without 10 μM NA functionally validating the exogenous expression of NAPRT **D.** Total NAD^+^ quantification post-24 hours in RH41 (NAPRT-silenced) and RH41^NAPRT+^ treated with respective IC_50_ concentration with or without 10 μM NA. **E-H.** CRISPR/Cas9-mediated knockout (KO) of NAPRT in endogenously NAPRT-expressing RH30 cells (RH30 NAPRT+). **E.** Protein expression of NAPRT, NAMPT, QPRT, and Vinculin in RH30 parentals and RH30 NAPRT-. **F-G.** Cell viability in RH30 models treated with increasing concentrations of OT-82 for 4 days with or without 10 μM NA functionally validating the KO of NAPRT **H.** Total NAD^+^ quantification post-24 hours in RH30 models treated with IC_50_ concentrations of OT-82 with or without 10 μM NA. The data is plotted as mean with error bars indicating SEM. ****p<0.0001; n.s., not significant.

### NAMPT-mediated NAD^+^ depletion leads to early-phase apoptosis in RMS cells

As we observed remarkable potency and decrease in cellular proliferation following NAMPTi in RMS cells with a loss of NAPRT, we sought out to decipher the mechanisms of cytotoxicity. Treatment with OT-82 induced only modest G2/M arrest following 48 hours of NAMPTi treatment in RH41 (NAPRT-silenced); however, NAMPTi induced little to no arrest in RH30 (NAPRT-expressing) without NA supplementation (**Figure 3A and B, Supplemental Figure S5A**). Prior studies in various tumor types have indicated that early phase apoptosis is observed following NAMPTi treatment (42). Most recently, in RMS, some models that exhibited full depletion of ATP following 72 to 120 hours of OT-82 treatment were shown to undergo necrosis (25). We treated RH41 and RH30 cells with OT-82 across multiple time-points and concentrations. After 48 hours, cleaved PARP1 signal was increased in both models across all concentrations of OT-82 (**Supplemental Figure S5B**). Next, we treated RH41 and RH30 cells with OT-82 and concurrent NA supplementation for 48 hours. Cells with intact NAPRT lost the cleaved-PARP1 signal with co-administration of NA (**Figure 3C**). To further corroborate this observation that apoptosis is ameliorated upon NA supplementation, we performed Annexin-V/propidium iodide (PI) staining. Supporting the immunoblot, in RH30 (NAPRT-expressing) cells, the induction of apoptosis was blunted when media was supplemented with NA (**Figure 3D-E; Supplemental Figure S5C**). Furthermore, in RH41 (NAPRT-silenced) cells, there was an increase in Annexin-V positive and double-positive (Annexin-V/PI positive) events indicating an induction of early to late apoptosis (**Figure 3F-G; Supplemental Figure S5C).** Collectively, these data demonstrate that RMS cell lines undergo apoptosis that may progress to necrosis due to intracellular metabolic dysregulation. Notably, these assays were performed at concentrations above the IC50 values obtained from 4–6 day viability studies to account for a shorter incubation period, which may explain differences in cell death mechanisms compared to prior reports (25). Importantly, cell death induced by NAMPT inhibition is rescued by NA co-administration in NAPRT-expressing models.

**Figure 3.**
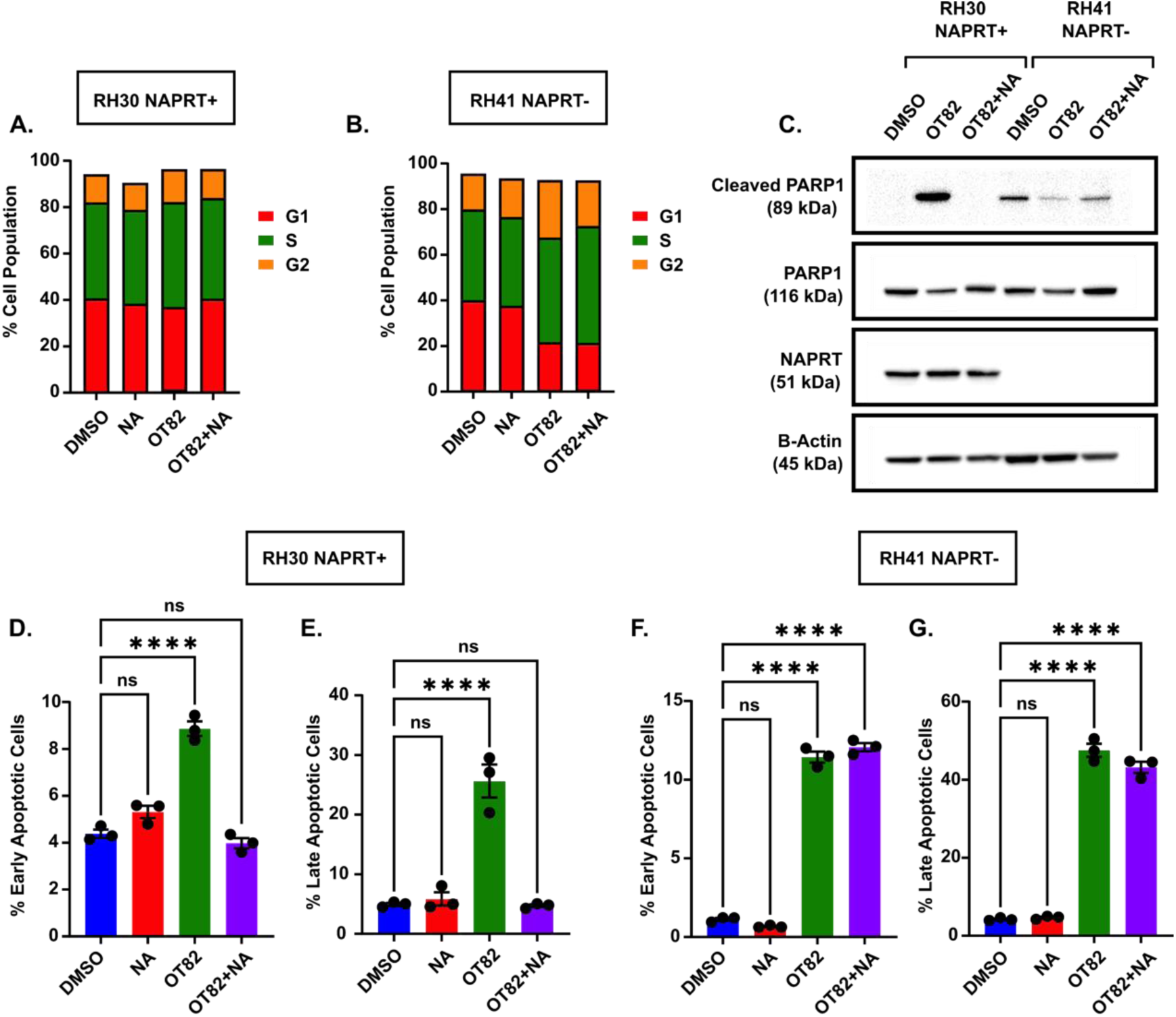
NAMPTi induces early to late-phase apoptosis that is reversible with NA in NAPRT-expressing RMS models. A-B. Cell cycle arrest after 48 hours of treatment with 0.1% DMSO, 10 μM NA, 10 nM OT-82, or the combination of OT-82 and NA in RH30 (NAPRT+) (**A**) and RH41 (NAPRT-) (**B**). **C.** Protein expression of Cleaved PARP1, PARP1, NAPRT, and Beta-actin on RH30 (NAPRT+) and RH41 (NAPRT-) treated with DMSO, 10 nM OT82, or combination of OT-82 and NA for 48 hours. **D-G**. Annexin-V/PI analysis after 48 hours of treatment of 0.1% DMSO, 10 uM NA, 10 nM OT-82, or the combination of OT-82 and NA on RH30 (NAPRT+) (**D-E**) and RH41 (NAPRT-) (**F-G**). The data represents mean with error bars as SEM. ****p<0.0001; n.s.,= not significant.

### NAMPTi demonstrates significant in vivo anti-tumor activity in RMS that is maintained with NA supplementation in NAPRT negative models

To assess the efficacy of NAMPTi and NA co-administration in the context of endogenous NAPRT expression *in vivo*, we treated orthotopic RMS models bearing RH30 tumors. Following a clinical schedule used in early-phase trials, OT-82 was administered daily for 3 days on/4 days off at 25 mg/kg (43). Furthermore, 25 mg/kg of OT-82 and 25 mg/kg of NA was co-administered in the supplementation cohort. In this endogenously NAPRT expressing model, tumors regressed following 2 cycles of treatment and survival was prolonged. However, we observed on-treatment progression after 3 cycles (**Figure 4A and D)**. Notably, NA co-administration completely abrogated anti-tumor activity with OT-82. Next, we performed *in-vivo* studies with RH41 NAPRT isogenic pairs. Tumor regression was observed immediately after 1 cycle of OT-82 treatment in the mice bearing RH41 parental (NAPRT-silenced) tumors **(Figure 4B).** Mice bearing RH41^NAPRT+^ tumors experienced tumor regression after 2 cycles of OT-82 treatment **(Figure 4C**). Survival was also improved in both RH41 parental and RH41^NAPRT+^ tumors **(Figure 4E** and **F).** Importantly, while NA co-administration did not diminish the efficacy of OT-82 in RH41 parental (NAPRT-silenced) tumors, RH41^NAPRT+^ tumors exhibited no response to treatment with NA co-administration (**Figure 4B,C,E,F**). We did not observe any obvious toxicity and mice maintained stable body weights in all three cohorts (**Figure 4G-I**).

**Figure 4.**
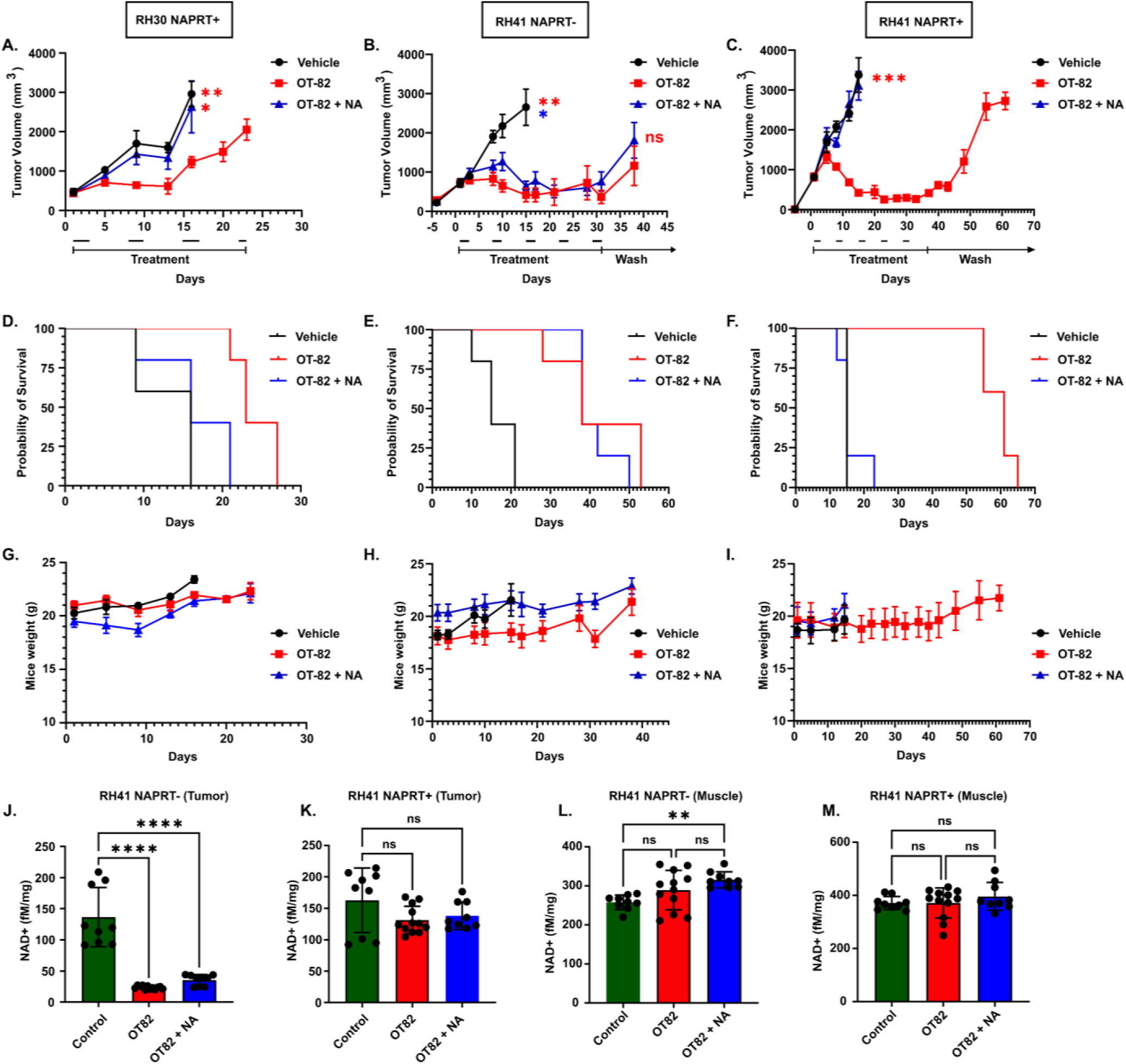
OT-82 leads to delayed tumor growth, prolonged survival and decreased NAD^+^ levels that is not rescued with NA in NAPRT-silenced xenograft models. A-F. Tumor growth curves (**A-C**), Kaplan-Meier plots (**D-F**), and body weight plots (**G-I**) of FOX SCID mice bearing RH30 (NAPRT+) (**A, D, G**), RH41 (NAPRT-) (**B, E, H**), RH41^NAPRT+^ (**C, F, I**) following treatment with 30% cyclodextrin vehicle, 25 mg/kg OT82, and 25 mg/kg NAMPTi in combination with 25 mg/kg NA. **J-K.** Total NAD^+^ levels in tumors harvested from RH41 (NAPRT-) (**J**) and RH41^NAPRT+^ (**K**) models treated with 1 cycle of 30% cyclodextrin vehicle, 25 mg/kg OT82, and 25 mg/kg NAMPTi in combination with 25 mg/kg NA. **L-M.** Total NAD^+^ levels in contralateral normal gastrocnemius muscle harvested from RH41 (NAPRT-) (**L**) and RH41^NAPRT+^ (**M**) treated with 1 cycle of 30% cyclodextrin vehicle, 25 mg/kg OT-82, and 25 mg/kg NAMPTi in combination with 25 mg/kg NA. For **J-M,** each treatment included 3–5 biological replicates, with 2 technical replicates per biological replicate. The data is plotted as mean with error bars indicating SEM. *p<0.05; **p<0.01; ***p<0.001; ****p<0.0001; n.s., not significant.

To assess the *in vivo* pharmacodynamic mechanism, total NAD^+^ in tumor samples was measured following 1 cycle of OT-82 with and without NA co-administration in the RH41 NAPRT isogenic model. In RH41 (NAPRT-silenced) tumors, NAD levels were significantly depleted after OT-82 treatment compared to vehicle control, even with exogenous NA administration (**Figure 4J**). Interestingly, in RH41^NAPRT+^ tumors, there was no statistically significant depletion of NAD^+^ after treatment regardless of NA administration even though anti-tumor efficacy was observed with prolonged treatment (**Figure 4K)**. Additionally, quantifying NAD^+^ pools in the contralateral normal muscle, there was no significant decrease in NAD^+^ in either of the OT-82 cohorts bearing either NAPRT+ or NAPRT-tumors (**Figure 4L** and **M**). To corroborate this finding, we analyzed tumor volumes after one treatment cycle, which entailed 3 days of treatment followed by 4 days of rest. RH41 (NAPRT-silenced) tumors demonstrated significant regression after 1 cycle of OT-82 alone or with NA supplementation. In contrast, no regression was observed in RH41^NAPRT+^ tumors after the same treatment cycle (**Supplemental Figure S6A**). After 2 cycles, a clear separation in tumor volumes between control and OT-82-treated groups was evident in both NAPRT-silenced and NAPRT-expressing tumors (**Supplemental Figure S6B**).

### NAPRT protein expression, methylation, and gene expression in RMS patient samples

To extend our findings in cell lines, we then evaluated for loss of NAPRT expression at the protein level in patient tumor samples via immunohistochemistry staining using a newly validated NAPRT monoclonal antibody (4A5D7)(9). First, we probed a pilot tissue microarray (TMA) generated at our institution consisting of 33 human RMS samples. Based on a semi-quantitative scoring system accounting for the percentage of positive tumor cells, 28% of RMS samples demonstrated loss of NAPRT protein expression classified as <1% of tumor cells showing any cytoplasmic or nuclear staining **(Figure 5A; Supplemental Table 2).** Next, we performed IHC staining in a TMA generated from RMS PDX models representing a total of 29 unique patients and again observed loss of NAPRT protein expression in roughly 30% of samples (**Figure 5B; Supplemental Table 3**). No significant difference in the frequency of NAPRT loss was observed across histologic subtypes or by FOXO1 fusion status in either the pilot patient cohort or the PDX TMA (**Figure 5C; Supplemental Figure 7A** and **B**). To corroborate these findings in a larger cohort of samples, we performed IHC staining in multiple validated TMAs previously generated from pediatric patients with RMS enrolled in COG clinical trials (**Supplemental Table 4).** The COG TMAs consisted of samples from a total of 109 patients 0 to 18-years-old. Using the same semi-quantitative scoring system utilized in the pilot TMA study, approximately 40% of pediatric RMS samples demonstrated loss of NAPRT protein expression **(Figure 5D; Supplemental Figure S7C).** Notably, NAPRT protein loss did not significantly differ by histologic subtype or FOXO1 fusion status (**Figure 5E** and **F**). Examination of clinical data from patients with total loss of NAPRT revealed a higher proportion of male patients, with a predominance of tumors arising in genitourinary sites (**Supplemental Figure S7D** and **E**). Furthermore, no significant associations were observed with tumor stage or metastatic status (**Supplemental Figure S7 F-K**).

**Figure 5.**
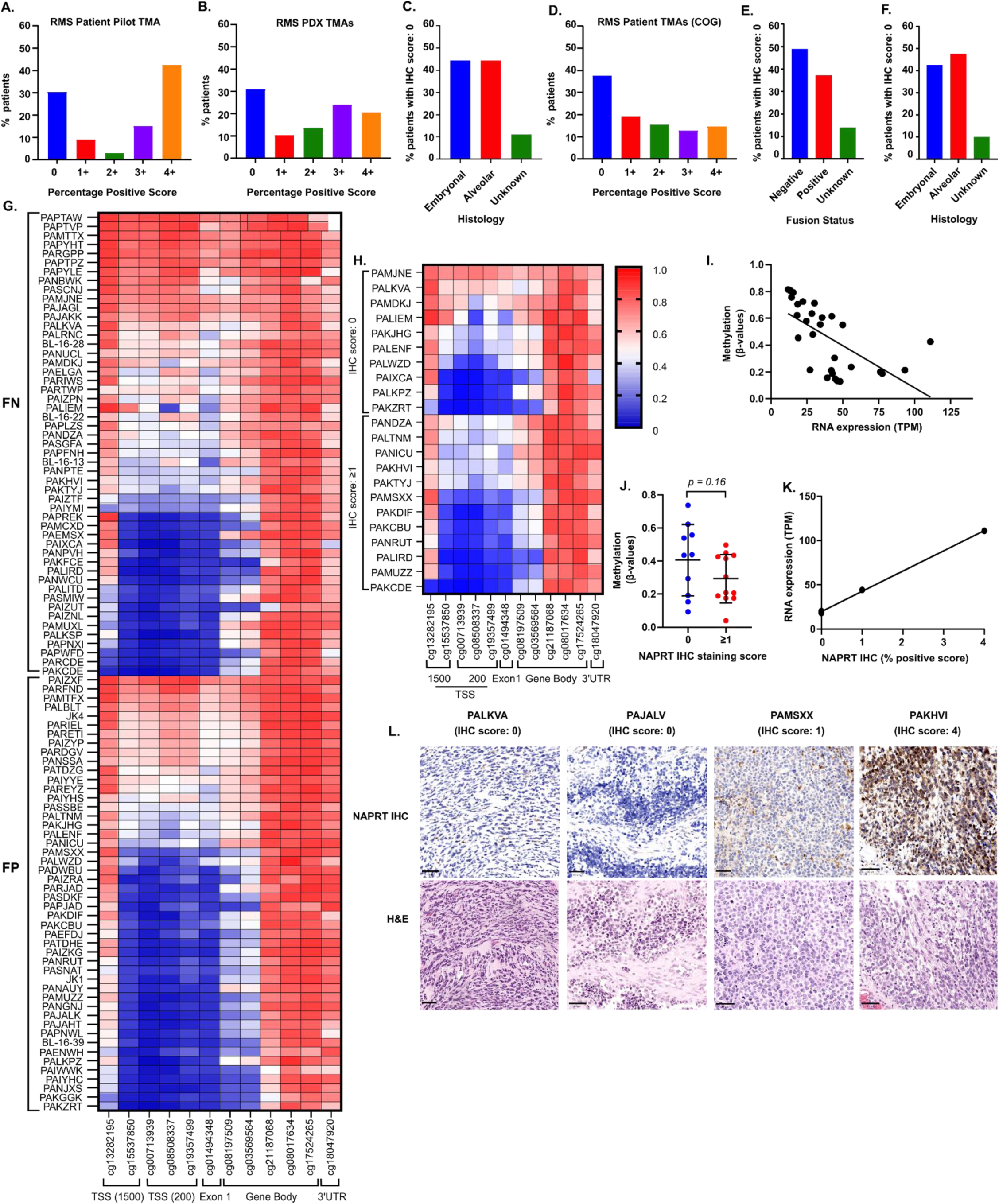
Loss of NAPRT expression in RMS patient samples. **A.** Summary of NAPRT IHC score in pilot tissue microarray (TMA) (n = 33 individual patients) using semi-quantitative scoring system based on percentage of positive tumor cells. **B-C.** Summary of NAPRT IHC score in patient derived xenograft (PDX) TMA (n = 29 individual RMS PDX) by percentage positive tumor cells (**B**) and by histology (**C**). **D-F.** Summary of NAPRT IHC score in Children’s Oncology Group (COG) TMAs (n = 109 individual patients) by percent positive tumor cells (**D**), fusion status (**E**), and histology (**F**). **G.** Heatmap representing methylation beta values of 12 CpG probes corresponding to the NAPRT1 gene (x-axis) of 99 pediatric RMS patients from COG TMA, stratified by fusion status (y-axis). **H.** Heatmap of 22 pediatric RMS patients with corresponding IHC and methylation data, stratified by total protein loss (top; IHC score = 0) or the presence of NAPRT protein (bottom; IHC score ≥1). **I.** Inverse relationship between RNA expression (TPM) and methylation beta values at the transcriptional start site (TSS) 200 and 1500 of 31 RMS patients (r^2^ = 0.38). **J.** Graph showing a negative correlation trend of beta values at TSS200 and TSS1500 regions with NAPRT IHC score for 22 patients in (**H**). **K.** Correlation between COG TMA NAPRT IHC score and RNA expression of four patients (r^2^ = 0.99). **L.** Representative NAPRT IHC (top) and hematoxylin and eosin (H&E; bottom) images in patients with corresponding IHC and transcriptomic data. In (**A, B, D**), percentage positive score: (Score = % Positive Cells) 0 = <1%; 1+ = 1-25%; 2+ = 25-50%; 3+ = 50-75%; 4+ = 75-100%.

Because NAPRT promoter methylation has been implicated as a driver of epigenetic silencing, we sought to correlate methylation with transcription and protein expression (9,13,26,28,44,45). We analyzed previously generated methylation data from 99 RMS patient samples through the Children’ Oncology Group (COG). Consistent with our findings in cell line models, we observed that a subset of RMS harbors hypermethylation of the NAPRT promotor across both FOXO1 fusion positive and negative tumors (**Figure 5G**). Of the COG samples with available methylation data, 31 had corresponding RNA-sequencing data and 22 had corresponding tissue in the TMA for IHC staining (**Figure 5H**). We observed an inverse correlation between the average methylation beta-values at the TSS200 and TSS1500 regions of the NAPRT promoter with gene expression, as measured by TPM (**Figure 5I)**. Interestingly, there was a trend towards higher average methylation beta-values at TSS regions in samples with complete loss of NAPRT protein staining as compared to those with any positive staining, although this did not reach statistical significance (**Figure 5J)**. Only 4 patients had corresponding transcriptomic and IHC data, which demonstrated a positive correlation between NAPRT gene expression and protein staining, despite the limited sample size (**Figure 5K** and **L**).

## Discussion

RMS remains a major therapeutic challenge, particularly in high-risk patients where there is a high unmet need for novel treatments to improve survival. Consistent with emerging evidence in pediatric bone and soft-tissue sarcomas, our findings demonstrate that RMS is highly vulnerable to NAMPT inhibition (23,25). Importantly, we build upon this foundational work by providing the first comprehensive analysis of NAPRT expression in pediatric RMS, leveraging patient-derived tumor samples, preclinical *in vivo* models, and isogenic cell line systems.

Probing methylation data within the DepMap, we found a high frequency of NAPRT gene promoter hypermethylation in RMS models. Our *in vitro* data show that NAPRT-deficient RMS cells are incapable of utilizing the Preiss-Handler pathway for NAD^+^ biosynthesis, and their survival is dependent on NAMPT activity. NAMPTi treatment led to profound cytotoxicity and NAD^+^ depletion in these cells, effects that were not reversed with nicotinic acid (NA) supplementation. In contrast, NAPRT-proficient RMS cells were rescued from NAMPTi-induced cytotoxicity with NA*. In vivo*, NAMPTi alone and with NA induced significant tumor regression in both endogenous and engineered NAPRT-deficient orthotopic RMS xenograft models. Notably, contrary to some prior reports, NA co-administration did not abrogate anti-tumor efficacy in NAPRT-deficient tumors but did preserve NAD^+^ levels and reversed anti-tumor efficacy in NAPRT-positive tissues (46). Interestingly, in RH41^NAPRT+^ tumors, there was no statistically significant depletion of NAD^+^ after 1 cycle of OT-82 treatment as compared to RH41 parental (NAPRT-silenced) tumors. This corresponded with the observation that RH41 parental tumors demonstrated significant regression after 1 cycle with or without NA supplementation, whereas no regression was observed in RH41^NAPRT+^ tumors at the same timepoint, even though robust anti-tumor efficacy was ultimately observed in both models with prolonged treatment. These data may suggest that NAPRT expression alters the kinetics of NAD^+^ depletion and cytotoxicity *in vivo*.

Based on these findings, our study supports the feasibility of a “protective rescue” paradigm, where co-treatment with NA could expand the therapeutic window of NAMPTi, potentially overcoming dose-limiting toxicities noted in preclinical and early-phase clinical trials (13,26,40,42). Based on the dose conversation between animals and humans (HED (mg / kg) = animal dose (mg / kg) × Km ratio (animal/human)), our dose of 25 mg/kg of NA supplementation for our *in vivo* studies in mice is approximately 75 mg/kg for humans, a dose well within the limits of safe daily NA intake (48). Importantly, dietary intake of NA has not been controlled for in previous clinical trials and may have contributed to lower-than-expected efficacy of NAMPTi by inadvertently rescuing NAPRT-expressing tumors. Prospective selection of NAPRT-silenced tumors could mitigate this confounder and improve therapeutic outcomes. Further pharmacokinetic and pharmacodynamic studies are warranted to characterize the NAD^+^ metabolome within the tumor microenvironment and to optimize NA dosing for future clinical translation of this therapeutic approach.

To further establish the translational relevance of NAPRT silencing in RMS, we evaluated promoter methylation, gene expression, and protein levels across multiple patient cohorts. Our findings demonstrate that a sizeable subset of RMS tumors exhibit NAPRT promoter hypermethylation, with approximately 30–40% of cases harboring loss of NAPRT protein expression. While promoter methylation is strongly implicated in NAPRT silencing, the modest correlation between methylation and protein loss observed in patient RMS tumors suggests additional regulatory mechanisms may be involved. These could include histone modifications, post-transcriptional regulation, or structural variants disrupting the NAPRT locus. Of note, our study is limited by the relatively small number of patient samples with matched protein, methylation, and transcriptomic data, restricting our ability to build a robust classifier for NAPRT status. Further investigation is warranted to determine the optimal cut-off of NAPRT expression and to refine biomarker-based patient selection strategies for future clinical trials. In addition, the heterogeneity of NAPRT expression within and between tumors warrants additional study, particularly in the context of potential clonal evolution under selective pressure from NAMPTi therapy.

NAPRT-silencing has also been reported in diffuse intrinsic pontine glioma and glioblastoma as well as fumarate hydratase deficient renal carcinoma due to varying epigenetic mechanisms (9,28). Evolutionarily, it is not well understood why some tumors prefer to rely on a specific NAD^+^ biosynthesis pathway and repress another. Interestingly, one study investigated the inherent loss of NAPRT through a tissue-lineage-based hypothesis, suggesting that tumors arising from normal tissues that do not depend on the Priess-Handler pathway, and therefore NAPRT, are more dependent on the NAMPT mediated salvage pathway (49). Intriguingly, skeletal muscle cells have been indicated as non-Priess-Handler pathway dependent (49). As a tumor that may arise from dysregulation in skeletal muscle differentiation, the tissue-lineage-based hypothesis may provide an explanation NAPRT silencing in RMS.

Furthermore, NAMPT plays an important role in mediating the downstream function of NAD⁺-dependent enzymes that influence multiple aspects of cancer biology. Among these, poly-ADP ribose polymerases (PARPs) are major consumers of NAD^+^ as it functions as an important substrate needed for PARP to form poly (ADP-ribose) chains, which recruit DNA damage response proteins. As such, previous studies have reported synergy between PARP inhibitors with NAMPTi (9,23,50). Moreover, co-treatment with temozolomide and NAMPTi has shown additive effects in preclinical glioma models, attributed a “hyper-vulnerability” secondary to NAD^+^ depletion from PARP1 activation (51). Exploring combination strategies that pair NAMPTi with DNA-damaging agents or DNA repair inhibitors represents an important future direction for clinical translation that bridges tumor metabolism and genomic instability.

Sirtuins (SIRT1-7) are another family of NAD⁺-dependent enzymes, and depletion of NAD^+^ has been shown to downregulate SIRT1 and, subsequently, upregulate PD-L1 expression (52). Additionally, NAD+ depletion has been shown to inactivate SIRT2 (53). Similarly, inactivation of SIRT2 following NAD⁺ loss has been linked to activation of the cGAS-STING pathway, suggesting that NAMPT inhibition may also have immunomodulatory consequences (54). Together, these findings highlight the broader biological implications of NAD⁺ depletion beyond cytotoxicity and support further investigation into how NAMPTi may reshape the tumor microenvironment through effects on DNA repair and immune signaling.

In conclusion, we identify NAPRT loss as a frequent and functionally relevant event in RMS that can be exploited therapeutically through NAD^+^ depletion with NAMPTi. Our work has important implications for the design of future clinical trials of NAMPT inhibitors in pediatric cancers. First, routine assessment of NAPRT status, via IHC or methylation analysis, could enable selection of patients most likely to benefit from NAMPTi monotherapy. Second, in patients with NAPRT-expressing tumors, NA co-administration may allow dose escalation while reducing off-target toxicities. Overall, these findings support further clinical study of NAMPTi in RMS via biomarker-driven clinical trials and rational combination therapies aimed at overcoming resistance and minimizing toxicity.

## Authors’ Contributions

Concept and design: A. Kim, P. Bhardwaj, J.C. Vasquez

Development of Methodology: A. Kim, P. Bhardwaj, S.J. Zhao, K.N. Lucas, K. J. Noronha, S. Friedman, J. Yi, J. Spurrier

Acquisition of data: A. Kim, P. Bhardwaj, S.J. Zhao, K.N. Lucas, K. J. Noronha, C.D. Heer, R. Morotti, R.K. Sundaram, S. Rehman, D. Bhatt, S. Ramakrishnan. W. Xue, D.A. Barkauskas, D. Hall, J.F. Shern, W. Sun, F. Barr, J. Yi, J. Spurrier, A. Yu, C.M. Heske,

Analysis and interpretation of data (e.g., statistical analysis, biostatistics, computational analysis): A. Kim, P. Bhardwaj, K.N. Lucas, S. Friedman, S. Ramakrishnan. W. Xue, D.A. Barkauskas, D. Hall, J.F. Shern, W. Sun, F. Barr, J. Yi

Writing, review, and/or revision of the manuscript: A. Kim, P. Bhardwaj, S.J. Zhao, C.M. Heske, J.C. Vasquez, C. Brenner

Administrative, technical, or material support: J.C. Vasquez, R.K. Sundaram, F.S. Delacruz, T. Feinberg Study supervision: J.C. Vasquez

## Acknowledgments

The authors would like to thank Dr. Ranjit S. Bindra and Dr. Douglas Spitz for support in conceptualization of the study and NAD^+^ metabolism expertise. The authors also thank Drs. Mitch Raponi and Marc Damelin from Alphina Therapeutics for providing anti-NAPRT antibodies for IHC staining and expertise on development of companion diagnostics. The authors also thank Dr. Corinne Linardic for providing the RH28 cell line. A. Kim is supported by Yale’s Cancer Biology Training Program (5T32CA193200-09). C.D. Heer is supported by the NIH/NCI award (K00CA245722). J.C. Vasquez is supported in part by the NIH/NCI award (1K08CA258796), Hyundai Hope on Wheels Scholar Hope Grant, the Robert Wood Johnson Harold Amos Medical Faculty Development Program, the Fund to Retain Clinical Scientists at Yale, sponsored by the Doris Duke Charitable Foundation award #2015216 and the Yale Center for Clinical Investigation, and by an American Cancer Society Institutional Research Grant, #IRG-21-132-60-IRG. This work was also supported by grants from the National Cancer Institute, National Institutes of Health to the Children’s Oncology Group (U10CA180886, U24CA196173, U10CA180899). J.F. Shern, F.G. Barr and C.M. Heske were supported by the Intramural Research Program of the National Cancer Institute. We thank the Yale Center for Genome Analysis and the Yale Keck Biotechnology Core for their assistance. The results shown here are in part based on data from the Cancer Dependency Map (https://depmap.org). We thank Yale Flow Cytometry for their assistance with their service. The Core is supported in part by an NCI Cancer Center Support Grant # NIH P30 CA016359. We thank the Yale Tissue Pathology Services Core for their assistance in generating tumor TMAs. The authors would like to acknowledge the Childhood Solid Tumor Network at St. Jude for providing PDX models. The content is solely the responsibility of the authors and does not necessarily represent the official views of the National Institutes of Health.

---

**Supplementary Figure 1. NAPRT-silenced RMS models are sensitive to FK866 and cannot be rescued with NA. A-B.** Cell viability in NAPRT-silenced RD (**A)** and RH41 (**B)** treated with increasing concentrations of FK866 for 6 days with or without 10 μM NA. **C-D.** Cell viability in NAPRT-expressing RH28 (**C**) and RH30 (**D)** treated with increasing concentrations of FK866 for 6 days with or without 10 μM NA. Total NAD^+^ quantification post-24 hours in RD (**E),** RH41 (**F**), RH28 (**G**), RH30 (**H**) treated with respective IC50 concentration of FK866 with or without 10 μM NA. **I.** NAPRT expression by western blot in PDX model (SJRHB13758). **J.** Cell viability in SJRHB13758 treated with increasing concentrations of OT-82 with and without 10 μM NA. The data is plotted as mean with error bars indicating SEM. ****p<0.0001; n.s., not significant.

**Supplementary Figure 2. NAPRT-deficient RMS models are not able to utilize NA to rescue cell viability A-C**. Relative cell viability of RH41 (NAPRT-silenced), treated with IC70 concentration of FK866 with or without 1 mM NMN **(A),** 10 μM NA (**B**), 100 μM NR (**C**). **D-F**. Relative cell viability of RH30 (NAPRT-expressing), treated with IC70 concentration of FK866 with or without 1 mM NMN **(D),** 10 μM NA (**E**), 100 μM NR (**F**). Points indicate mean and error bars indicate SEM. ****p<0.0001; n.s., not significant.

**Supplementary Figure 3. Generation of plasmid based NAPRT overexpressing isogenic model. A-C.** Flow-sorted single cell clones of endogenously NAPRT-silenced RH41 parental cells transfected to exogenously express NAPRT (RH41 NAPRT+). **A.** Protein expression of NAPRT in RH41 parental and clones 1-4. **B-C.** Cell viability in RH41 NAPRT+ clones treated with increasing concentrations of FK866 **(B)** and OT-82 **(C)** for 4 days with or without 10 μM NA functionally validating the exogenous expression of NAPRT. **D-E.** Pooled population of endogenously NAPRT-silenced RD transfected to exogenously express NAPRT (RD NAPRT+). **D.** Protein expression of NAPRT in RD parental and RD NAPRT+. **E.** Cell viability in RD NAPRT+ pooled population treated with increasing concentrations of FK866 for 4 days with or without 10 μM NA functionally validating the exogenous expression of NAPRT. Points indicate mean and error bars indicate SEM.

**Supplementary Figure 4. Functional validation of single-cell clones of CRISPR/Cas9 ribonucleoprotein (RNP) based NAPRT knock-out model. A-C.** Expression and functional validation of CRISPR/Cas9 RNP based KO of NAPRT utilizing 2 sgRNA constructs; CRISPR948669_SGM (gRNA1) and CRISPR948672_SGM (gRNA2). **A.** Protein expression of NAPRT and Beta-actin in RH30 parentals (Parentals) and RH30 NAPRT-single cell clones. **B-C.** Cell viability in RH30 NAPRT-clones from gRNA1 (**B**) and gRNA2 (**C**) treated with increasing concentrations of OT-82 for 4 days with or without 10 μM NA, functionally validating the exogenous KO of NAPRT. Points indicate mean and error bars indicate SEM.

**Supplementary Figure 5. Induction of apoptosis in RMS models with OT-82 treatment. A.** Histogram showing RH41 (NAPRT-) and RH30 (NAPRT+) cell cycles after 48 hours of treatment with 0.1% DMSO, 10 uM NA, 10 nM OT-82, or the combination of OT-82 and NA. **B.** Protein expression of PARP1, Cleaved PARP1, NAPRT, and Beta-actin on RH30 (NAPRT+) and RH41WT (NAPRT-) treated with 0.1% DMSO, 5 nM, 10 nM, 20 nM, and 30 nM of OT-82 for 48 hours. **C**. Representative Annexin-V/PI flow plots (left) and quantification (right) after 48 hours of treatment of 0.1% DMSO, 10 uM NA, 10 nM OT-82, or the combination of OT-82 and NA on RH30 (NAPRT+) and RH41 (NAPRT-). The data is plotted as mean with error bars indicating SEM. *p<0.05; **p<0.01; ***p<0.001; n.s.,= not significant.

**Supplementary** Figure 6**. Tumor volumes in RMS isogenic models after OT-82 treatment. A.** Dot plots for tumor volumes after 1 cycle (3 days on and 4 days off) of control (30% cyclodextrin), 25 mg/kg OT-82, and 25 mg/kg OT-82 with NA co-administration on FOX SCID mice bearing RH41 parental (NAPRT-; left) and RH41^NAPRT+^ overexpression (NAPRT+; right) orthotopic tumors. **B.** Dot plots for tumor volumes after 2 cycles of control, OT-82, and OT-82 with NA co-administration. * p<0.05; **p<0.01; ****p<0.0001; n.s., not significant.

**Supplementary Figure 7. Clinical Correlates of NAPRT Expression in RMS patient samples. A-B.** Pilot TMA NAPRT IHC score by histology (**A**) and fusion status (**B**). **C.** Representative NAPRT IHC (top) and hematoxylin and eosin (H&E; bottom) images in patient tumors from COG TMA with range of IHC semi-quantitative score based on percentage positive tumor cells. **D-G.** Clinical annotations of patients with IHC score of 0 (total loss of NAPRT) by sex (**D**), site of primary tumor (**E**), stage (**F**), and metastatic vs non-metastatic disease (**G**). **H-K.** Clinical annotations of patients with IHC score of ≥1 by sex (**H**), site of primary tumor (**I**), stage (**J**), and metastatic vs non-metastatic disease (**K**). **L.** Dot plot of beta-values at probes within TSS200 and TSS1500 regions by fusion status in 99 patients. Error bar indicates SD. *p<0.05. **M.** Correlation between NAPRT IHC score and NAPRT beta-values at CpG probes present within TSS200 and TSS1500 in 22 patients.

**Supplemental Table S1.** List of antibodies, company and catalog number, and dilution utilized for western blots.

|  |  |  |
| --- | --- | --- |
| Vinculin | Cell signaling technology 13901 | 1:1000 |
| GAPDH | Protein Tech, 6004-1-Ig | 1:1000 |
| NAPRT | Proteintech 66159-1-Ig | 1:3000 |
| NAMPT | Cell signaling technology 86634 | 1:1000 |
| QPRT | Abcam 171944 | 1:1000 |
| Beta-actin | Cell signaling technology 3700 | 1:1000 |
| Cleaved PARP1 | Cell signaling technology 5625 | 1:1000 |
| PARP1 | Cell signaling technology 9542 | 1:1000 |

**Supplemental Table S2.** Summary of pilot TMA.

|  |  |
| --- | --- |
| Individual patients | 33 |
| Median age yrs (range) | 18.5 (0-37) |
| Total cores | 58 |
| Single core | 11 |
| Replicate cores | 47 |
| Fusion Status of cores |  |
| Fusion Negative | 10 |
| Fusion Positive | 14 |
| Other | 34 |

**Supplemental Table S3.** Summary of PDX TMA.

|  |  |
| --- | --- |
| Individual patients | 29 |
| Histologic subtype |  |
| Alveolar | 15 |
| Embryonal | 12 |
| MyoD1 mutant | 2 |
| Total cores | 55 |
| Single core | 3 |
| Replicate cores | 52 |

**Supplemental Table S4.** Summary of Children’s Oncology Group TMA.

| **Supplemental Table S4.** Summary of Children's Oncology Group TMA. |  |
| --- | --- |
| Individual patients | 109 |
| Median age yrs (range) | 9 (0-18) |
| Total cores | 278 |
| Single core | 35 |
| Replicate cores | 243 |
| Fusion status of cores |  |
| Fusion Negative | 51 |
| Fusion Positive | 41 |
| Other | 17 |

