## Supplemental figures for "NAPRT expression and epigenetic regulation in pediatric rhabdomyosarcoma as a potential biomarker for NAMPT inhibition"

### Supplementary figure 1

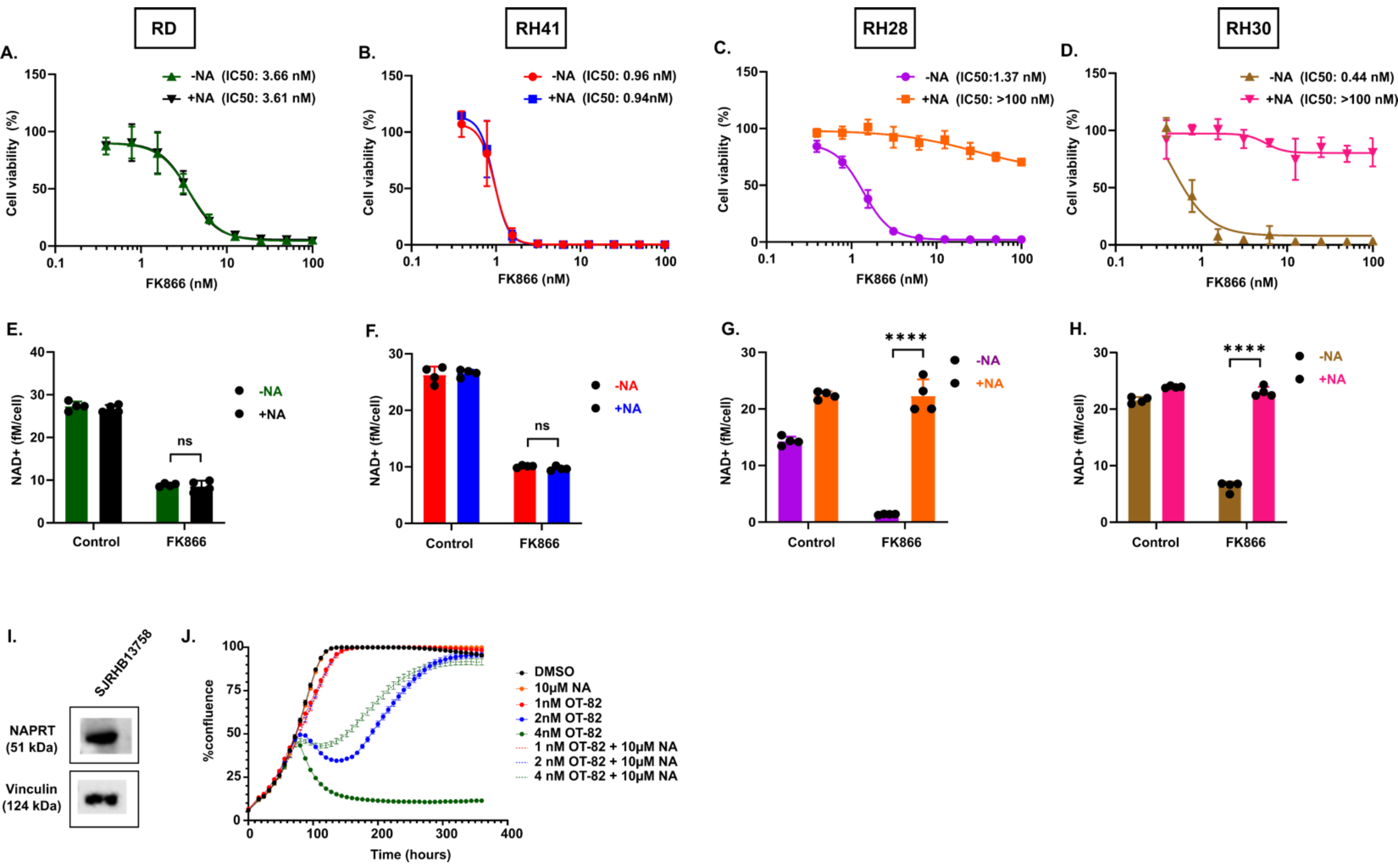

### Supplementary figure 2

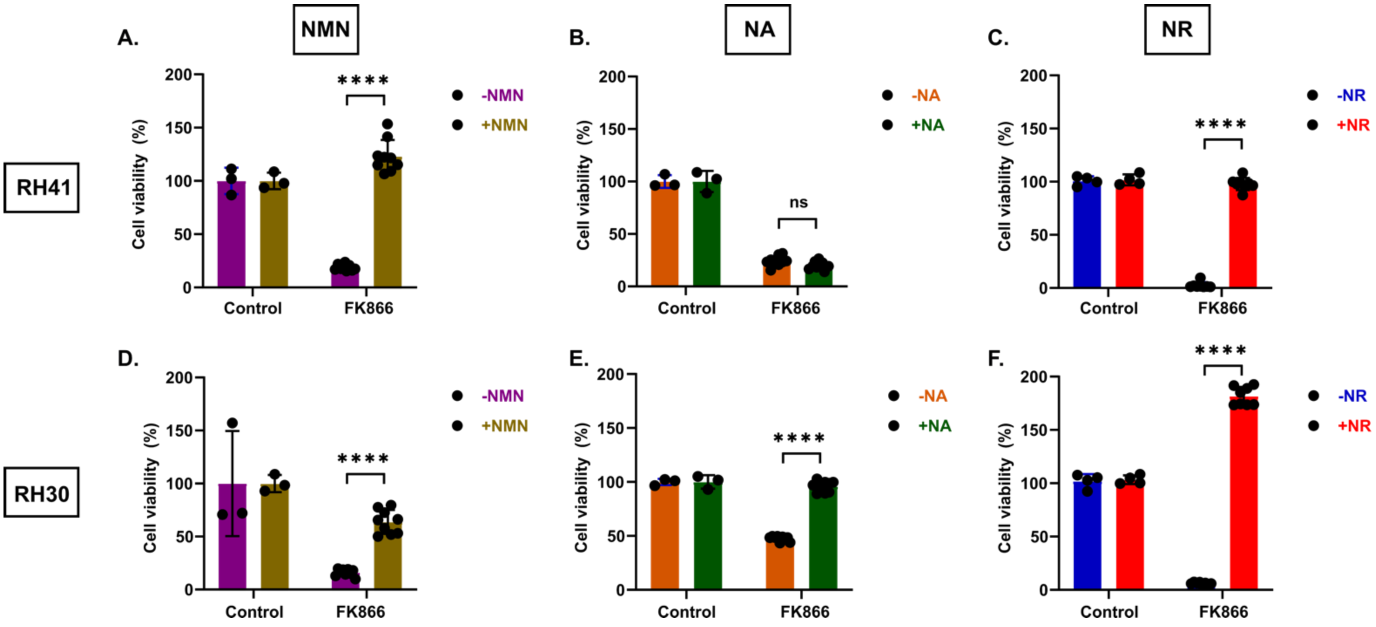

### Supplementary figure 3

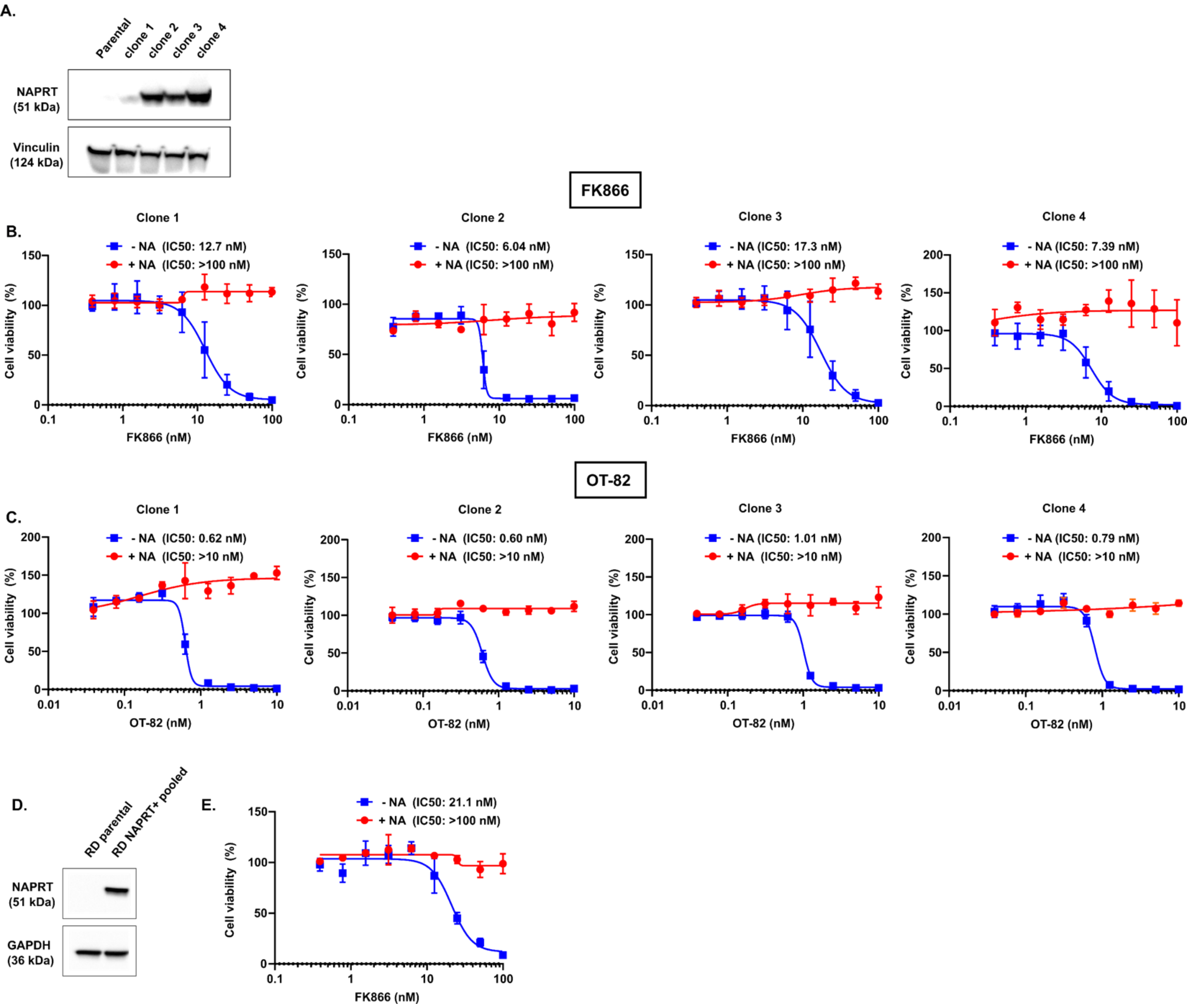

### Supplementary figure 4

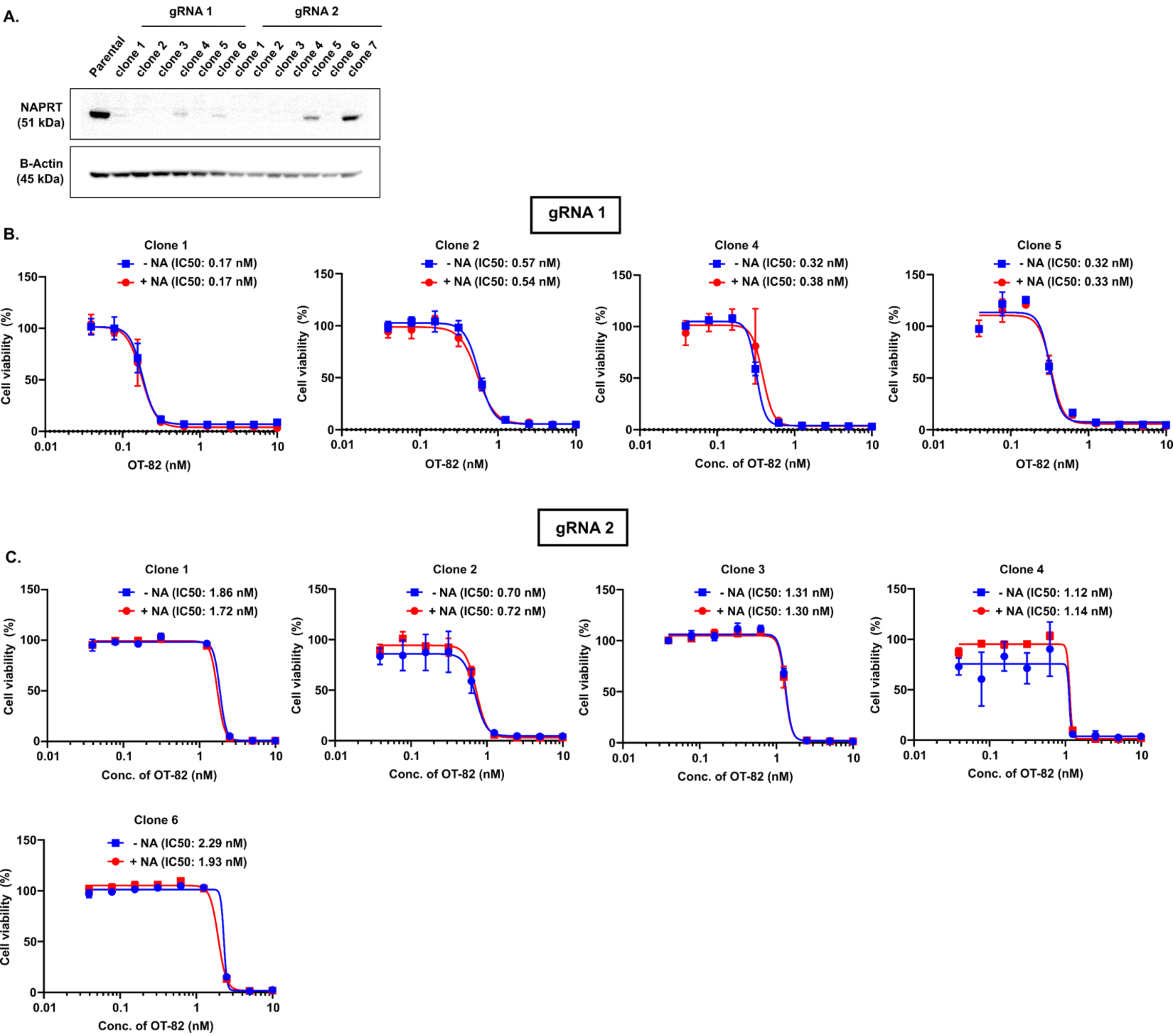

### Supplementary figure 5

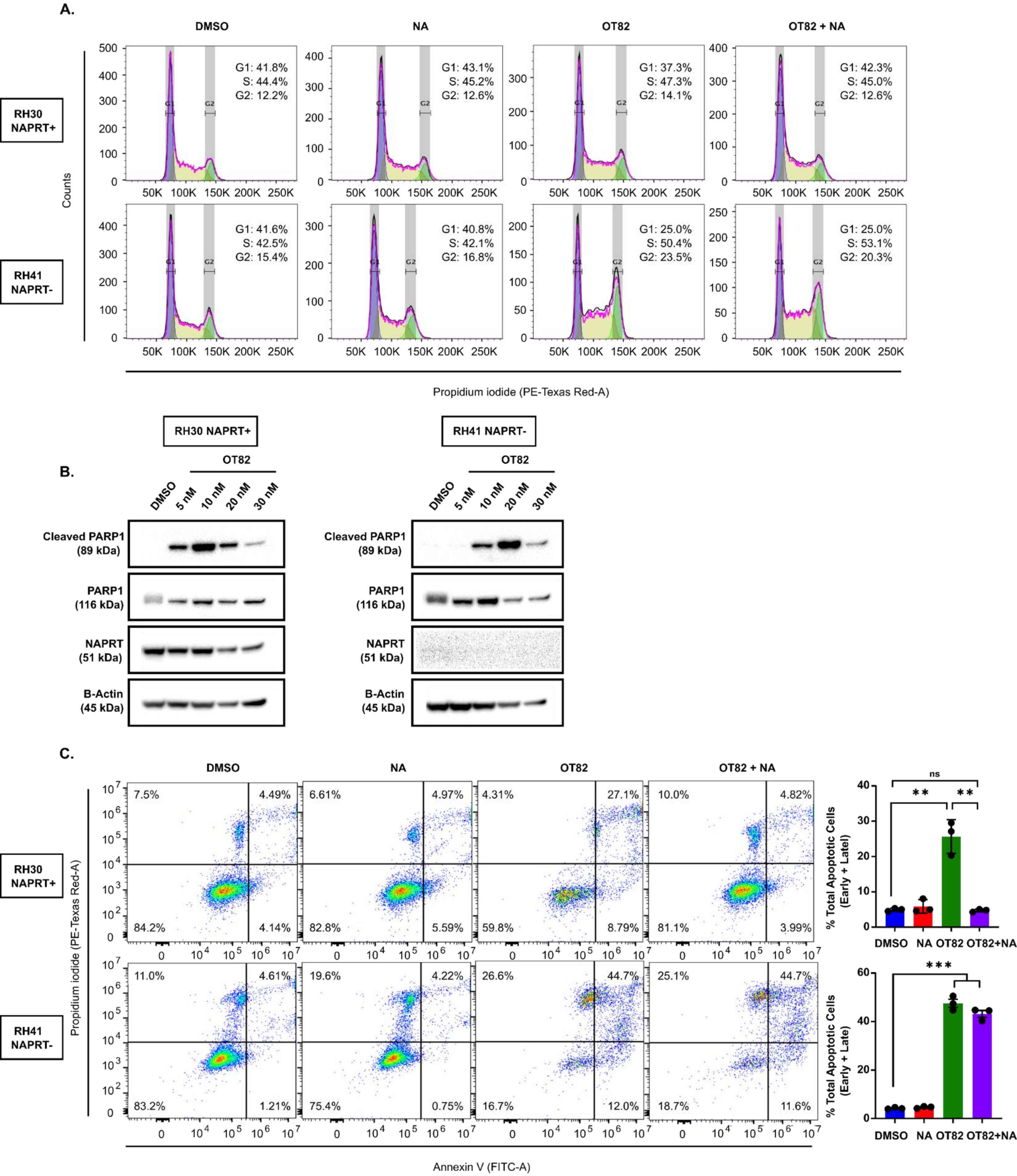

### Supplementary figure 6

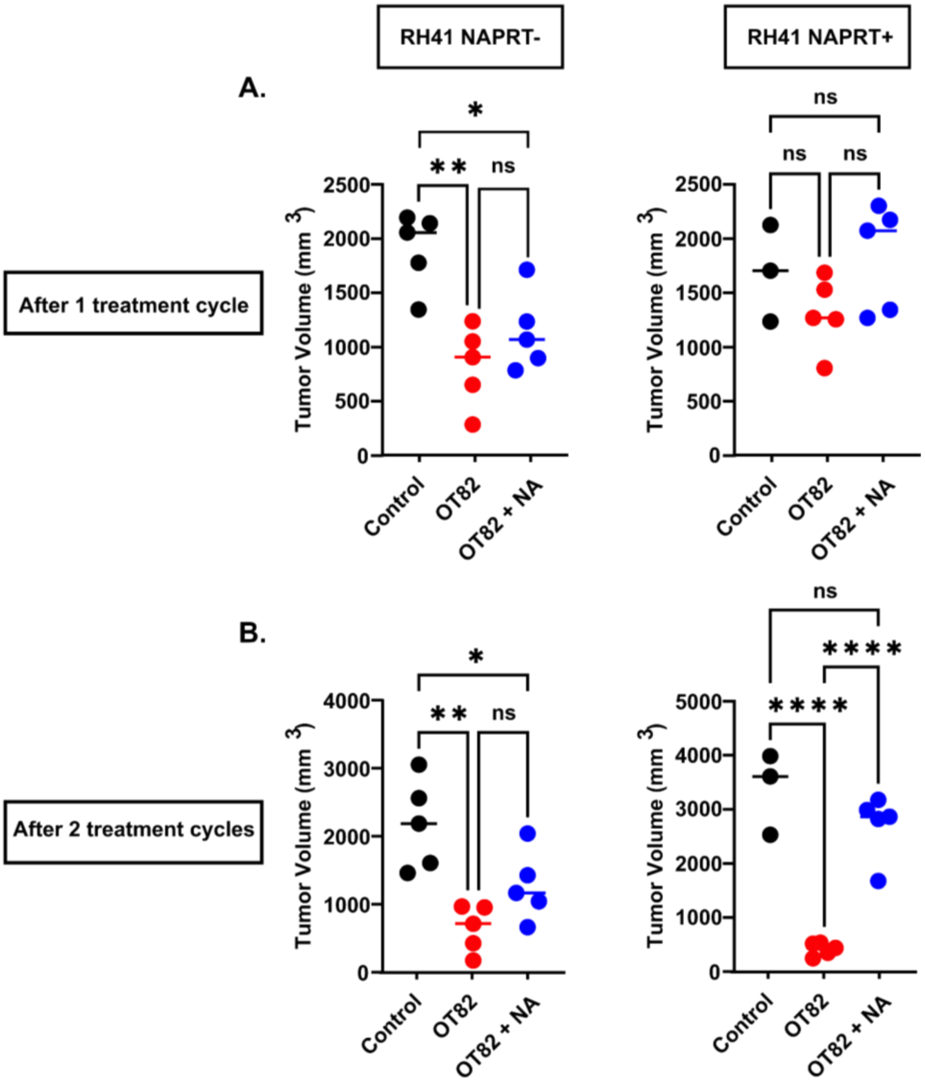

### Supplementary figure 7

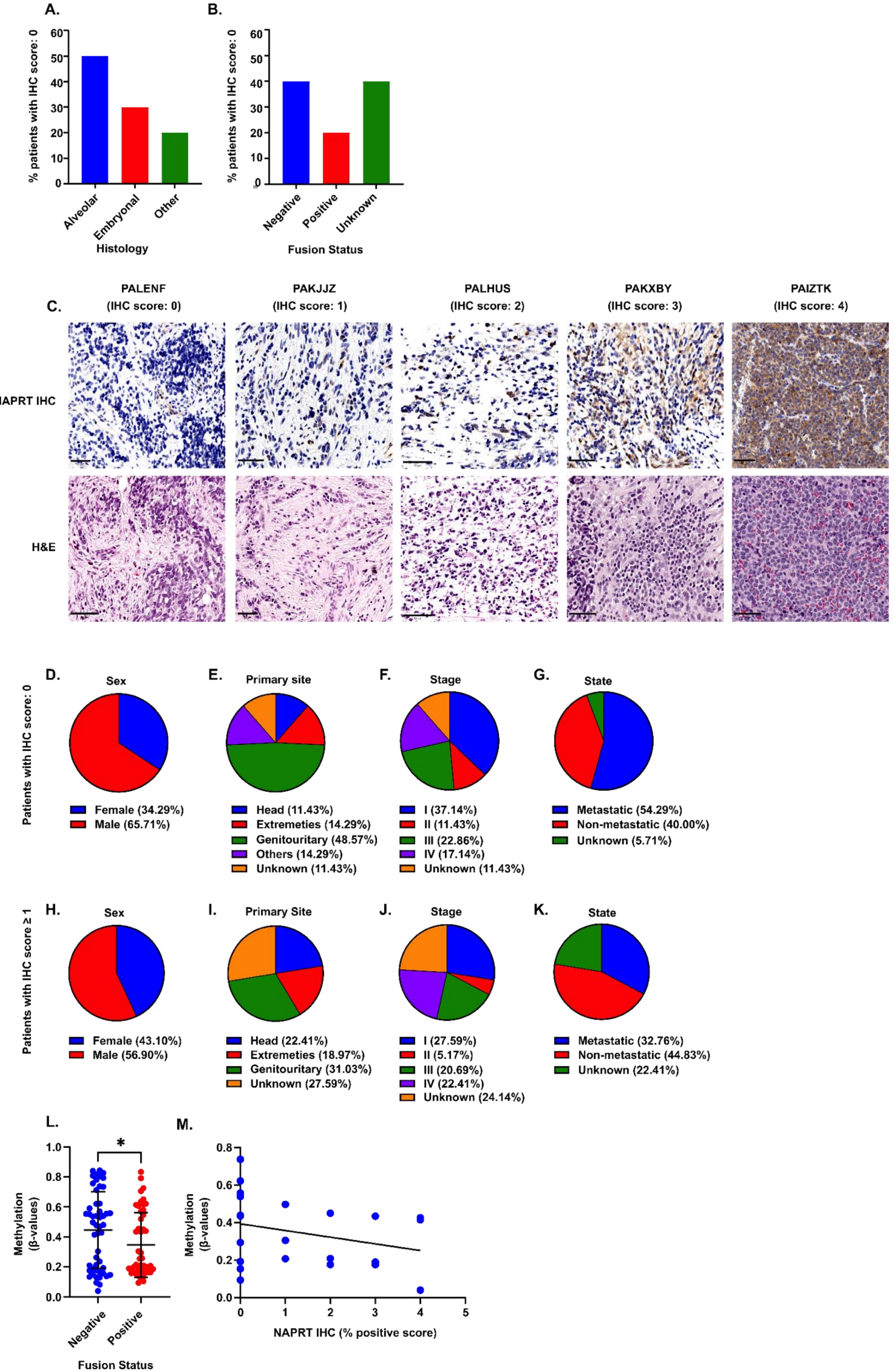
